# Honey bee functional genomics using symbiont-mediated RNAi

**DOI:** 10.1101/2022.04.22.489157

**Authors:** Patrick J. Lariviere, Sean P. Leonard, Richard D. Horak, J. Elijah Powell, Jeffrey E. Barrick

**Affiliations:** Department of Molecular Biosciences, The University of Texas at Austin, Austin, TX 78712, USA; Department of Integrative Biology, The University of Texas at Austin, Austin, TX 78712, USA

## Abstract

Bees are indispensable pollinators and model organisms for studying social behavior, development, and cognition. However, their eusociality makes it difficult to use standard forward genetic approaches to study gene function. To fill this gap, we engineered the bee gut bacterium *Snodgrassella alvi* to induce a host RNA interference response that reduces expression of a targeted gene. To employ this FUGUES (**<u>FU</u>**nctional **<u>G</u>**enomics **<u>U</u>**sing **<u>E</u>**ngineered **<u>S</u>**ymbionts) procedure, a double-stranded RNA expression plasmid is cloned in *Escherichia coli* using Golden Gate assembly and then transferred to *S. alvi*. Adult worker bees are then colonized with engineered *S. alvi*. Finally, gene knockdown is verified through qRT-PCR, and bee phenotypes of interest can be further assessed. Expression of targeted genes is reduced by as much as 50-75% throughout the entire bee body by five days after colonization. This protocol can be accomplished in four weeks by bee researchers with microbiology and molecular cloning skills.

## Introduction

The western honey bee, *Apis mellifera*, supports global agriculture and is a model organism for studies of insect cognition, sociality, and gut microbiomes. Despite this widespread use and economic importance, genetic tools for bees are underdeveloped. Most studies to date have examined differences in phenotypes due to natural genetic variation or how gene expression changes after an experimental treatment^1–4^.While these studies can suggest candidate genes involved in different biological processes and pathways, they are correlative and cannot rigorously establish that changes in gene expression cause a change in behavior or other phenotypes. Disrupting or lowering the expression of a target gene and observing the corresponding phenotypic effects is a fundamental approach for establishing gene function.

RNA interference (RNAi) is a highly conserved antiviral immune system that has been exploited to suppress gene expression and investigate gene function in many organisms, including insects, nematodes, and plants^5–8^. It responds to the presence of double-stranded RNA (dsRNA) in cells. Researchers commonly induce RNAi by injecting or feeding purified dsRNA or dsRNA-producing *Escherichia coli* cells^8^. Persistent induction of RNAi has also been achieved in insect pest species by engineering natural gut symbionts to stably express dsRNA^9^.

We recently showed that symbiont-mediated RNAi can be used to robustly, persistently, and systemically knock down gene expression in bees. We refer to this approach as FUGUES, for **<u>fu</u>**nctional **<u>g</u>**enomics **<u>u</u>**sing **<u>e</u>**ngineered **<u>s</u>**ymbionts. Here we provide a protocol for performing FUGUES to establish connections between genes and phenotypes in adult worker honey bees.

### Development of the protocol

The discovery of a core community of gut bacteria in honey bees^10–12^ laid the groundwork for the development of FUGUES. This community is remarkably conserved and stable throughout a worker honey bee’s adult life. One species of particular importance, *Snodgrassella alvi*, closely associates with the gut lining^12^ and induces host immune responses that regulate microbiome composition^13^. Shortly after the discovery of these core gut bacteria, Kwong *et al*. axenically cultured some of these species, including *S. alvi*^14^. They additionally showed that *S. alvi* could be re-administered to honey bees to achieve colonization with laboratory isolates^15^.

We subsequently developed the bee gut microbiome toolkit (BTK) for genetically modifying multiple species of bee gut bacteria^16^. The key components of the BTK include promoters that function in diverse bacteria and a broad-host-range plasmid that is compatible with using conjugation to deliver DNA to bacteria lacking established transformation protocols. These universal plasmid components allow the BTK to be used for genetically modifying other bacteria, including insect symbionts such as *Serratia symbiotica* from aphids^17^.

Recently, we used these approaches for culturing and genetically manipulating symbionts to engineer *S. alvi* to alter bee gene expression and protect bees from pathogens^18^. We reasoned that *S. alvi* was a suitable platform for affecting host biology because (1) *S. alvi* colonizes bees exceptionally well, even from a starting inoculum of only a few cells^18^, and (2) *S. alvi* lives in the bee gut in close association with host tissues as a biofilm lining the ileum^10,12^. These characteristics may facilitate exchange of bioactive molecules between *S. alvi* and its bee host.

We previously showed three applications of *S. alvi* as a platform for symbiont-mediated RNAi^18^. First, we demonstrated honey bee FUGUES by knocking down expression of an insulin receptor, increasing the sensitivity of bees to sucrose and leading to weight gain. Knockdown of the target gene was measured 5 days after colonization and persisted through 10 days after colonization.

We also showed that expression of dsRNA sequences targeting other organisms could be used to improve bee survival following exposure to a viral pathogen and to kill mites that parasitize bees.

### Applications of the method

Symbiont-mediated RNAi in honey bees can be used for functional genomics and pathogen protection. By knocking down genes and discovering their functions using FUGUES, researchers may also uncover new ways of regulating bee gene expression to benefit their health or improve pollination efficacy. These methods are intended for laboratory experiments and must be used only for research under approved institutional biosafety and biocontainment protocols. Further testing will be needed to establish whether genetically modified *S. alvi* can be used safely in normal apiculture where colonized bees would mix with other bees in the environment.

The FUGUES approach described here can potentially be applied to study other bee species, particularly those with *S. alvi* or closely related *Snodgrassella* species in their core microbiomes. Two additional *Apis* species (*A. andreniformis* and *A. cerana*) as well as many or all bumble bee species (*Bombus* spp.) also have abundant amounts of *Snodgrassella* in their microbiomes^19–22^. Additionally, certain stingless bees contain *Snodgrassella*, including *Partamona helleri, Trigona* spp. (*T. hyalinata* and *T. spinipes*), and *Tetragonula* spp. (*T. davenporti, T. pagdeni*, and possibly *T. carbonaria*)^20,23^. Both bumble bees and stingless bees are ecologically important natural pollinators, and the former are also used by commercial pollination services^23,24^. Therefore, these groups of insects are attractive targets for symbiont-mediated RNAi using our platform. Because *Snodgrassella* strains tend to be better at colonizing the bee species with which they have co-evolved^15,20^, different strains may be more effective for *Bombus* and stingless bee applications. Engineering other *S. alvi* strains to express dsRNA using the approach detailed here is expected to induce a similar RNAi response in other bees.

Beyond bees, symbiont-mediated RNAi has been described in three other insect species – kissing bugs (*Rhodnius prolixus*), western flower thrips (*Frankliniella occidentalis*), and diamondback moths (*Plutella xylostella*) – indicating wider potential applications^9,25–27^. FUGUES leverages a toolkit for assembling broad-host-range plasmids with an RSF1010 origin of replication that is compatible with both Gram-positive and Gram-negative bacteria^16,28^. Therefore, the dsRNA-expression plasmids described here are expected to function in many bacterial species, including other symbionts. More broadly, our hope is that this protocol can serve as a resource and template for groups interested in establishing symbiont-mediated RNAi and FUGUES in other insect and animal species for which suitable bacterial symbionts can be identified.

### Comparison with other methods

To date, multiple gene knockdown and knockout strategies have been reported in honey bees. Transposon mutagenesis in honey bees has been described in two studies^29–31^. However, it is difficult to successfully employ because it requires laborious injection techniques and achieves only a low mutagenesis efficiency, so it is not widely used. More recently, CRISPR/Cas genome editing has been used to generate either homozygous mutant drones or heterozygous mutant females in a number of studies^29,32–39^. However, CRISPR/Cas-based knockouts also require technically demanding injections^35^. Very few gene expression profiles have been quantified in mutant females, making this promising technology difficult to fully assess at this time^32^.

RNAi is the mostly widely used method for knocking down gene expression in bees. Though it is effective and reliable in diverse invertebrate species, including *C. elegans* and various insect model systems, RNAi has had varying degrees of success in attenuating gene expression in honey bees^29,40,41^. Successful gene knockdown by RNAi can be variable and depend on multiple factors, including target-specific effects (strength of target gene expression and tissue being targeted), dsRNA sequence design, dsRNA dosage, and method of dsRNA administration^29,40,41^. As this protocol describes a novel method of dsRNA delivery, we will focus on the state of the art of this aspect of RNAi in bees. To date, the majority of studies implementing RNAi in bees have involved either dsRNA injection or feeding^41^.

Injecting purified dsRNA is commonly used to knock down genes in *A. mellifera*^35,41^. This approach requires expression and purification of dsRNA from an *in vitro* system or *E. coli*^41,42^. dsRNA injection leads to variable knockdown depending on the gene. Results have ranged from a strong >70-75% decrease in mRNA levels for *Vg* and *amGPdh* to a weaker ∼30-40% decrease in mRNA levels for *AmOA1* or the hypopharyngeal amylase gene, for example^29,40,43–48^. Despite its wide usage, dsRNA injection does pose a few major drawbacks. First, it requires a high level of technical skill to perform, and the manual process of injecting individual bees presents a major bottleneck that limits throughput^49^. Second, gene knockdown by dsRNA injection is necessarily transient, as the injected dsRNA has a finite half-life^49^. Finally, the injection process is invasive and can cause physical trauma to the bee, potentially initiating stress responses and complicating the interpretation of downstream assays^41,49^.

Feeding purified dsRNA has also been used to induce RNAi in *A. mellifera*^41^. Like injection, this approach requires expression and purification of dsRNA. In contrast to injection, large numbers of bees and even whole hives can be fed dsRNA, as demonstrated in studies that suppress bee pathogens such as Israeli acute paralysis virus and *Varroa* mites^50–52^. dsRNA feeding has also been used to knock down bee genes, though – as with injection – the effectiveness varies depending on the gene^29,40,41,46,53–58^. Another downside of this approach is that it requires continuous addition of dsRNA to the bees’ feed to sustain gene knockdown.

The FUGUES approach of using symbionts to deliver dsRNA has key advantages compared to conventional methods of administering dsRNA via feeding or injection. First, FUGUES leads to both sustained and robust gene knockdown, because dsRNA is produced *in situ* and is continuously transferred to the host. Second, to our knowledge, FUGUES offers the highest scalability of any gene knockdown approach in bees, since a large batch of bees can be readily inoculated with a small amount of engineered *S. alvi*. Finally, administration of engineered *S. alvi* to bees is simple, eliminating the technical expertise required for dsRNA injection. In short, FUGUES leverages the power of the symbiont to more readily facilitate gene knockdown in bees compared to existing methods.

### Overview of the procedure

We provide a complete protocol for implementing symbiont-mediated RNAi in honey bees for FUGUES (Fig. 1). First, a dsRNA expression vector targeting an *A. mellifera* gene of interest is designed, assembled, and transformed into an *E. coli* cloning strain. Next, the dsRNA expression vector is transferred to the bee symbiont *S. alvi*. Subsequently, honey bees are inoculated with the engineered symbiont. Finally, knockdown of gene expression is validated using qRT-PCR on RNA isolated from bee tissues.

**Figure 1.**
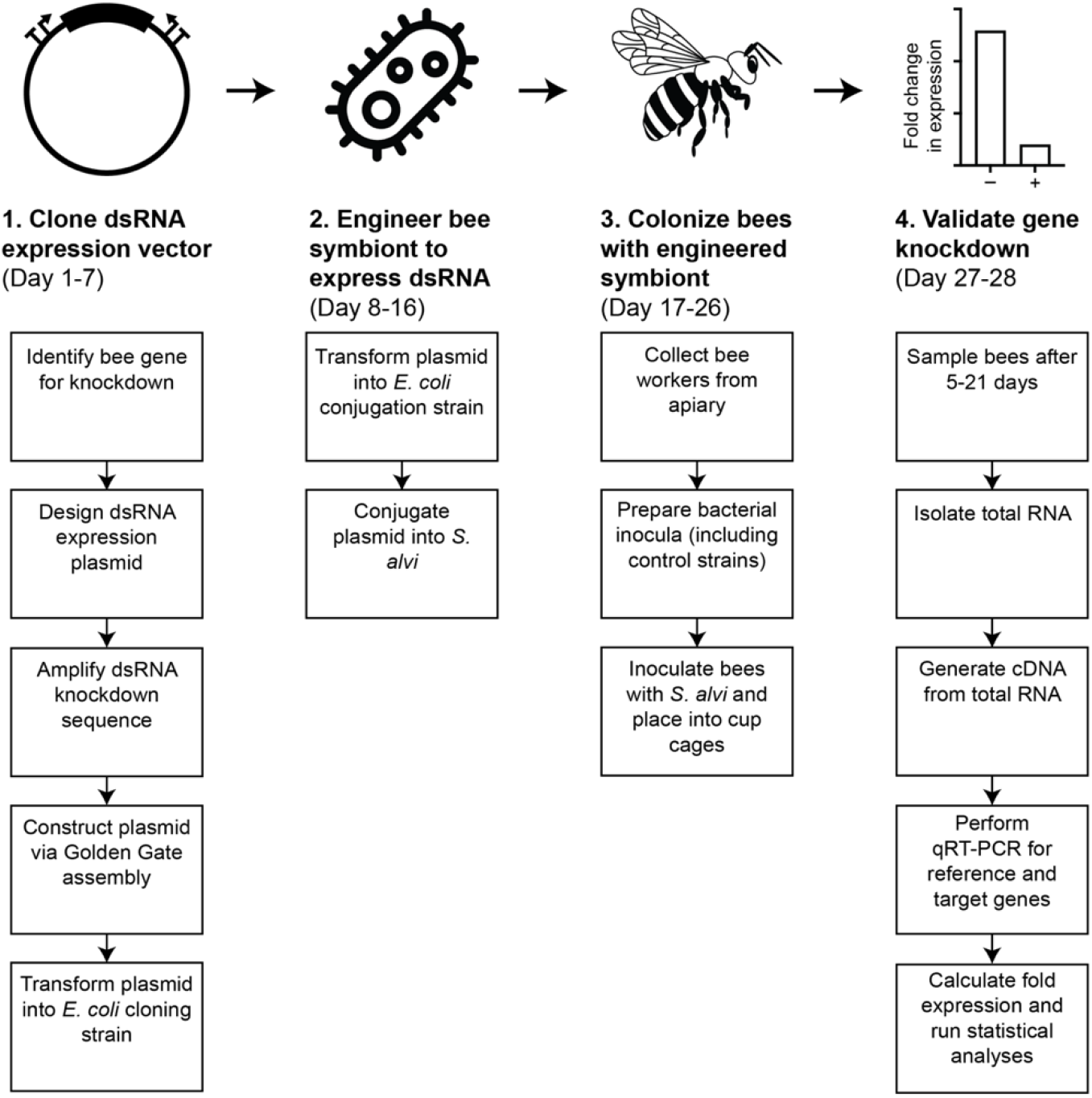
Workflow for using symbiont-mediated RNAi to study gene function in bees. Schematic of FUGUES workflow. In phase 1 (cloning) the dsRNA expression vector is designed, constructed using Golden Gate assembly, and transformed into *E. coli*. In phase 2 (transfer to bee symbiont), the dsRNA expression vector is conjugated into *S. alvi*. In phase 3 (bee inoculation), bees are collected and inoculated with engineered *S. alvi*. In phase 4 (gene knockdown validation), total RNA is collected from bees, reverse transcribed into cDNA, then used as template for qRT-PCR reactions using reference and target gene primers. Fold expression is then calculated to quantify changes in gene expression. The procedure as described takes a total of approximately 28 days, including 16 days to design, build, and validate the plasmid and engineered symbiont strains, 10 days to colonize and then house the honey bees, and 2 days to validate the knockdown. While we describe the process for a single target gene, multiple knockout targets can be designed, built, and tested in parallel.

### Experimental design

Here, we provide more detailed overviews of each of the four main phases of FUGUES. In addition to providing specific information regarding materials and methods, we pay special attention to controls and decision points where a researcher will need to adjust parameters to suit their study. Important controls include comparing the results to experiments in which *S. alvi* expresses an off-target dsRNA, to distinguish effects resulting from knockdown of a targeted bee gene from nonspecific immune responses^18^. This protocol can be adapted to any bee gene with a sufficient length that a dsRNA can be designed that matches part of it and there is enough remaining sequence for designing qRT-PCR primers for monitoring knockdown.

#### Clone dsRNA expression vector (Steps 1-16)

The first phase of FUGUES is to construct a dsRNA expression vector targeting a honey bee gene for knockdown. While deciding which gene should be chosen for knockdown will ultimately depend on the question a researcher is trying to answer, we provide a few general guidelines here. First, remember that FUGUES will only affect gene expression during the adult phase of a bee’s life. The functions of genes at earlier developmental phases cannot be studied using this approach. Second, check for paralogs of the gene of interest. If these exist, consider designing constructs to test knockdown of each one. Third, check to see if there are multiple transcript isoforms of the gene of interest. If these are found, perform an alignment to generate a consensus sequence that targets all of them or a specific subset for knockdown.

Once a gene has been selected, a piece of it is cloned into a dsRNA expression plasmid. To do this, a dsRNA knockdown region from the coding sequence of the gene of interest, typically 400 bp or longer, is chosen, and primers are designed to amplify this region. BsaI cut sites are appended to the forward and reverse primers for assembly with the plasmid backbone. We provide sequences for primers used to amplify portions of the *inR1* and *def1* genes, which can be used as positive controls. Knockdown of *inR1* was previously reported^18^, and knockdown of *def1* is reported for the first time here. At this stage, qPCR primers should also be designed to amplify a ∼100 bp region of the coding sequence. These will be used later in the protocol for validating gene knockdown. qPCR primers must amplify a region outside of the dsRNA knockdown region, to prevent them from detecting dsRNA produced by *S. alvi*. We provide sequences for *inR1* and *def1* qPCR primers that can be used to validate that gene expression is being knocked down using the positive controls.

Once primers have been obtained, the dsRNA knockdown region is amplified with PCR. The template used for PCR will depend on the design of the dsRNA knockdown region: honey bee cDNA can be used for all constructs, and honey bee genomic DNA can be used for constructs contained within a single exon. Following PCR, amplicons are purified, and cloned into a dsRNA expression vector. The assembled plasmid is transformed into *E. coli* cells and plated. The following day, three non-fluorescent colonies are picked and used for plasmid miniprep. Proper insertion of the dsRNA construct into the plasmid is validated by sequencing.

#### Engineer bee symbiont to express dsRNA (Steps 17-31)

After the dsRNA expression vector has been made, it is transferred to the bee symbiont *S. alvi*. To do this, the dsRNA expression vector must first be transformed into an auxotrophic, conjugation-competent *E. coli* strain, such as MFD*pir* (which is auxotrophic for diaminopimelic acid, DAP). Other auxotrophic conjugation-competent *E. coli* strains could also be used, such as ST18, a 5-aminolevulinic acid auxotroph^59^. Electrocompetent stocks of MFD*pir* are made and electroporated with the dsRNA expression vector. Transformants are plated on media containing spectinomycin and DAP following recovery. The following day, colonies are picked and saved as glycerol stocks.

To initiate conjugation, *S. alvi* and MFD*pir* cells are grown, then pelleted, and resuspended in PBS. Cells are combined, then plated on media containing DAP, to allow for conjugation to occur. The next day, cells are scraped, washed, and plated on media containing spectinomycin (but no DAP), to select for cells that have taken up the plasmid and to select against MFD*pir*. Transconjugant colonies are passaged onto a second plate containing spectinomycin. After 2-3 days of growth, glycerol stocks are made from a final scape of these plates.

The plasmids pNR and pDS-GFP serve as negative controls for gene knockdown and off-target RNAi responses. They must be transferred into *S. alvi* if this is the first time a researcher is using this protocol. These plasmids or the pDS-Def1 and pDS-InR1 plasmids that can used as positive controls for demonstrating knockdown of targeted genes via FUGUES can also be used as positive controls for the electroporation and conjugation steps.

#### Colonize bees with engineered symbiont (Steps 32-42)

Once *S. alvi* is engineered to express dsRNA, it is ready to use to colonize bees. The engineered symbiont is struck out onto an agar plate. For each experiment, engineered *S. alvi* strains are used from three groups: the experimental group (*S. alvi* containing a plasmid targeting a gene of interest), the positive control group (*S. alvi* containing pDS-Def1 or pDS-InR1), and the negative control group (*S. alvi* containing pDS-GFP). The use of the positive control group is optional but highly recommended for new users of this protocol. Twenty-four hours prior to bee colonization, engineered *S. alvi* is passaged onto a fresh agar plate. Serially passaging *S. alvi* is not recommended, as lab-adapted symbionts may be less effective at colonizing bees.

Bees should now be obtained for colonization through one of two methods: (1) allowing young adult bees to naturally emerge on their own from the frame or (2) pulling pupae and allowing bees to finish development in plastic brood chambers. Bees that emerge on their own are easier to obtain in large numbers but may also have environmental bacteria present in their guts. Bees obtained by pulling pupae are nearly germ-free, but this procedure is more laborious and requires more technical expertise. The timing of these methods differs, so it is important to decide which method will be used early in the process of planning an experiment. Both methods are described in step 35, with the method in which bees emerge on their own presented as the default.

When bees have been obtained for colonization, the plate containing engineered *S. alvi* is scraped and washed. The *S. alvi* suspension is diluted, mixed with a bee feeding solution, and used to coat 20-30 bees per condition. Colonized bees are transferred to cup cages and, after a brief fast, are given sucrose feeding solution supplemented with spectinomycin. Bees are maintained for one week, with the feed and solution being replaced every three days.

#### Validate gene knockdown (Steps 43-56)

After one week, bee samples are collected and gene knockdown is validated using qPCR on bee cDNA. For each condition and timepoint, approximately ten bees are removed from cup cages and frozen. Depending on where the gene of interest is expressed, RNA for assessing knockdown can be isolated from entire bees or dissected heads, abdomens, guts, or other body parts. Extracted RNA is then copied into cDNA through reverse transcription. qPCR primers for the gene of interest and reference gene are then tested for adequate efficiency. Subsequently, qPCR is performed on bee cDNA derived from the experimental group (gene of interest knocked down) and control groups (Positive control: *inR1* or *def1* knockdown; Negative control: *gfp* mock knockdown), using primers for the gene of interest and multiple reference genes (such as *rps18* and *gapdh*)^60,61^. Ct values determined by qPCR are used to calculate ΔΔCt, which is used to determine the relative expression of the gene of interest (compared to the reference gene) for the experimental and control groups^62,63^. Gene expression data are visualized with graphing software and statistical analyses are performed to determine whether a significant reduction in expression of the target gene has been achieved.

Depending on the desired outcome, bees can be assessed for phenotypes of interest or gene expression changes resulting from the dsRNA treatment as early as 3-5 days after colonization. Multiple time points can be analyzed to assess the persistence of gene knockdown over time. We have tested that knockdown persists through least 10 days after colonization^18^ and do not have any evidence that it disappears or is attenuated at any point later in the lives of bees.

### Expertise needed to implement the protocol

This protocol assumes that the reader has access to an apiary and experience working with honey bees. We provide instructions that relate to special considerations for obtaining uncolonized bees and administering engineered bacteria to them. However, the details of honey bee husbandry and maintenance are outside the scope of this protocol. All microbiology, cloning, conjugation, and molecular biology experiments can be performed in a standard molecular microbiology laboratory with access to a qPCR machine and a CO_2_ incubator for culturing *S. alvi*.

### Limitations

Some limitations of the current iteration of honey bee FUGUES should be considered before a researcher begins this protocol. FUGUES is only suitable for use in adult worker bees, as larvae and pupae do not harbor the gut symbiont used to deliver dsRNA. As a result, genes involved in early development at the larval and pupal stages cannot be studied. Because FUGUES constitutively induces the host RNAi response beginning early in an adult bee’s life, it may perturb normal host physiology in ways that complicate the study of genes that function at later life stages, when bees transition to different worker roles, for example. Additionally, FUGUES has not been tested in queen bees, which have different gut microbiomes from workers. We also caution that biosafety and biocontainment protocols approved by an investigator’s institution must be put into place before using this procedure, as with any genetically modified organisms. Because honey bee hives are typically maintained outdoors in open environments that do not meet these requirements, in-hive behaviors cannot be tested without access to specialized containment facilities. Finally, because an antibiotic is required to maintain the engineered plasmid/symbiont, the bee microbiome is not in its natural state during a FUGUES experiment. Some of these limitations could be addressed in the future by engineering bee symbiont species that colonize larvae, integrating dsRNA constructs into the bacterial chromosome, or using inducible promoters that allow for temporal control of dsRNA expression.

### Reagents

#### Clone dsRNA expression vector

Plasmid pBTK800 (Addgene #179209) or plasmids pBTK403, pBTK301, pYTK002, pYTK072, pBTK150, pBTK151 (Addgene #110599, #110593, #65109, #65179, #183126, #183127)

DNeasy Blood & Tissue Kit (QIAGEN, #69504)

Quick-RNA Tissue/Insect Kit (Zymo, #R2030)

1^st^ strand cDNA Synthesis Kit (Takara, #6110A)

Q5 Hot Start High-fidelity 2X Master Mix (NEB, #M0494S)

QIAquick PCR purification kit (QIAGEN, #28104)

5-alpha Competent *E. coli*, High Efficiency (NEB, #C2987H)

Golden Gate Assembly Kit, BsaI-HFv2 (NEB, #E1601S)

Quick Plasmid Miniprep Kit (PureLink, #K210010)

LB broth and agar plates

SOC medium

Spectinomycin (Spectinomycin dihydrochloride pentahydrate) (CAS #22189-32-8)

Chloramphenicol (CAS #56-75-7)

Glycerol (CAS #56-81-5)

Molecular grade water (DNase-free and RNase-free) (CAS #7732-18-5)

Primers designed for cloning dsRNA region for gene of interest (See step #2)

#### Engineer bee symbiont to express dsRNA

*Snodgrassella alvi* strain wkB2 (ATCC #BAA-2449)

Negative control plasmids pNR and pDS-GFP (Addgene #183131 and #183129)

Positive control plasmids pDS-Def1 and pDS-InR1 (optional)

(Addgene #183125 and #183130)

MFD*pir* or similar auxotrophic *E. coli* conjugation strain (e.g., ST18, DSM 22074) 2,6-Diaminopimelic acid (DAP) (CAS #583-93-7)

5-aminolevulinic acid (if using ST18 instead of MFD) (Sigma-Aldrich, CAS #5451-09-2) Columbia Blood Agar Base (Hardy-Diagnostics, #C5451)

Sterile Sheep’s Blood (Lampire, 7239001-1LTR)

Phosphate-Buffered Saline (PBS)

#### Colonize bees with engineered symbiont

Sucrose, (CAS #57-50-1)

Pollen (Prairie River Honey Farm, Bulk Fresh Bee Pollen Pure Raw Natural Nebraska

Bee Pollen 6 lbs; Pollen shipped by user to Sadex Corporation for irradiation)

#### Validate gene knockdown

Quick-RNA Tissue/Insect Kit (Zymo, #R2030)

1^st^ strand cDNA Synthesis Kit (Takara, #6110A)

DNase I Set (Zymo, #E1010)

RNA Clean and Concentrator-25 (Zymo, #R1017)

OneTaq 2X Master Mix with Standard Buffer (NEB, #M0482S)

SYBR Green PCR Master Mix (Thermo-Fisher, #4309155)

qPCR primers designed for gene of interest (See step 3)

qPCR primers for reference gene (See step 3)

### Equipment

#### Clone dsRNA expression vector

Computer with DNA analysis software (Benchling, Geneious, or other)

Shaking incubator (New Brunswick I24Benchtop Incubator Shaker)

Benchtop centrifuge with rotor for 1.5 mL tubes (Eppendorf Centrifuge 5430)

Thermocycler (Eppendorf Mastercycler Nexus Gradient)

Agarose gel electrophoresis system (Galileo Bioscience Minigel System)

Mini-Beadbeater-96 cell disruptor (BioSpec Products, #1001)

Molecular biology grade consumables (Pipette tips, pipette, tubes, racks)

Cryovials (Thermo Scientific, #37518)

Nanodrop (Thermo Scientific NanoDrop Lite Spectrophotometer)

Benchtop spectrophotometer that can measure OD_600_ (Eppendorf BioPhotometer, #6131)

Disposable cuvettes (Thermo Fisher, #14955127)

#### Engineer bee symbiont to express dsRNA

Electroporator (Bio-Rad Gene Pulser Xcell Electroporation System)

Electroporation cuvettes (1 mm) (Thermo Fisher, #FB101)

CO_2_ incubator (Panasonic CO_2_ Incubator, #KM-CC17RU1A)

Microbiological loops, 1 µL (Biologix, #65-001)

#### Colonize bees with engineered symbiont

Standard honey bee husbandry equipment (hives, bee suit, smoker, smoker fuel, hive tool)

Custom built frame cage

Cup cage materials

- Plastic cups (Solo, #TP10D)
- Petri dish lids (Corning, #BP93B-102)
- Filter paper (Thermo Fisher, #09-795C)
- 96 well plate for pollen trough (Greiner, #655101)
- Tape (Fisherbrand, #15901R)
- 10 ml tubes for cup cage feeders (Sarstedt, #62.9924.283)

Environmental chamber with humidity control for rearing bees in lab (Percival Incubator, #I36NL)

50 mL conical tubes (Corning, #430829)

Vacuum filter bottles (Nalgene, #567-0020)

Ice bucket

#### Validate gene knockdown

qPCR machine (Eppendorf Mastercycler ep realplex Real-time PCR System)

Computer for qPCR machine

96 or 384 well qPCR plates (Bio-Rad HSL9645)

Optical quality plate covers (Excel scientific, #09-795C)

Multichannel pipettes and tips

### Reagent setup

#### Prepare sucrose solution (50% [w/w] sucrose:water) for honey bees

1. Mix 8 lb sucrose with 1 gallon hot tap water, then filter sterilize.

#### Prepare LB media and agar plates

1. Prepare media according to manufacturer’s instructions.
2. Prepare selective LB agar plates by adding spectinomycin to a final concentration of 60 µg/mL.
3. Prepare selective LB agar plates by adding chloramphenicol to a final concentration of 20 µg/mL.

#### Prepare DAP stock solution for supplementing media

1. Add 342.4 mg of DAP to 30 mL of H_2_O to create a 60 mM stock solution.
2. Shake at 37 °C for ∼30 min to fully dissolve, then filter sterilize.
3. Store aliquots frozen at −20 °C.
4. This is a 200× stock. Use at a final concentration of 0.3 mM in media.

#### Prepare Columbia Blood Agar Plates

1. Combine 44 g Columbia media into 950 mL sterile H_2_O with a stir bar.
2. Autoclave at 121 °C for ≥ 20 minutes.
3. Remove media from autoclave and let cool to < 60 °C.
4. Add 50 mL sterile sheep’s blood.
5. For selective plates, add spectinomycin to a final concentration of 30 µg/mL to molten agar with blood.
6. Pour agar into sterile petri plates and let cool overnight.
7. Store agar plates at 4 °C prior to use.
8. Depending on humidity, plates should be dried before use by leaving uncovered under a sterile laminar flow hood for 30 minutes prior to use.

#### Prepare honey bee cup cages

1. Poke airholes into side of plastic cups (Fig. 4C) and hole for feeding tube into bottom of cup (Fig. 4C).
2. Cut 96 well plate into pollen troughs. In the fume hood, use a flame heated paring knife to cut grooves into a 96 well plate. Then snap the plate into 2×3 well pollen troughs.
3. Place filter paper onto petri dish lid.
4. Place pollen trough on the petri dish lid and fill with irradiated pollen.
5. Place cup (with holes) onto the petri dish lid and tape to secure.
6. Place piece of tape over hole on top of the cup if not using feeding tube, to prevent bees from escaping.

### Procedure

#### Clone dsRNA expression vector

##### Identify bee gene for knockdown

1. To begin the steps needed for cloning the dsRNA expression vector (Fig. 2A), either download the latest honey bee genome assembly (Amel_HAv3.1, GCF_003254395.2 as of 10/31/2021) from NCBI and find the target gene locus using genome viewing software, such as Geneious (paid license required, https://www.geneious.com) or locate and show the sequence of the target gene locus using the online NCBI Genome Data Viewer (available for free use, https://www.ncbi.nlm.nih.gov/genome/gdv/browser/genome/?id=GCF_003254395.2). (10 minutes)

**Figure 2.**
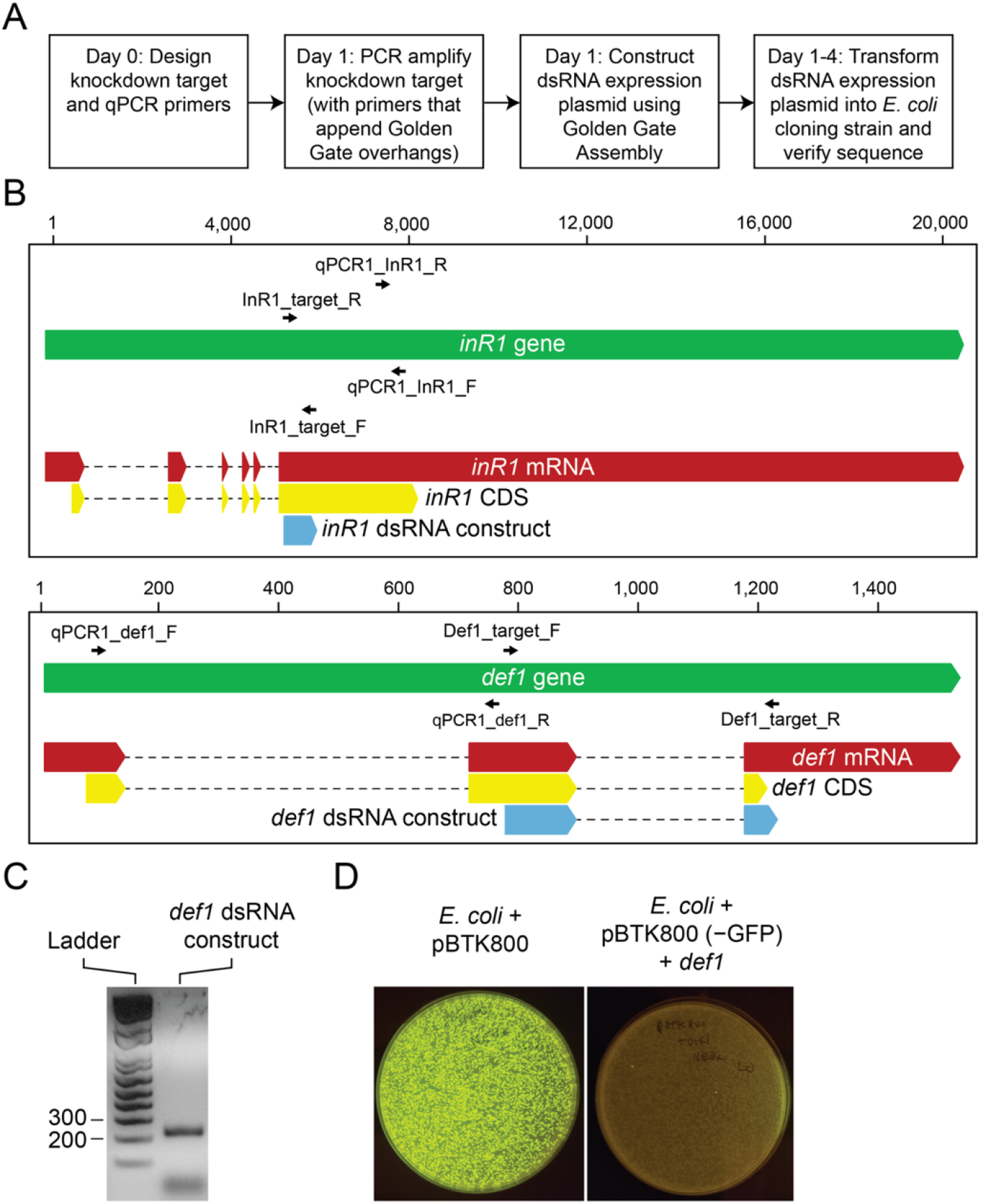
dsRNA expression vector cloning. A. Overview of the dsRNA expression vector cloning workflow. B. Maps of example genes chosen for knockdown: *inR1* (top) and *def1* (bottom). Each map displays the architecture of the gene (green), mRNA (red), coding sequence (CDS) (yellow), and dsRNA construct PCR product (blue). Primer sites for amplifying the knockdown target for *inR1* and *def1* and qPCR primers for both genes are displayed as small arrows. The horizontal scale (in nucleotides) is shown above each gene map. The *inR1* gene is pictured in the reverse orientation to how it is encoded in the *A. mellifera* chromosome, so its R and F primers appear flipped here. Also note that PCR is performed on honey bee cDNA in these examples, so only the regions present in the mRNA (red) will be amplified. Information for *inR1* is from a published study^18^. C. PCR amplification of the *def1* dsRNA target sequence. The amplicon runs at the expected 230 bp size on a 2% agarose gel (200 and 300 bp ladder standards indicated). *D. def1* dsRNA expression plasmid assembly. pBTK800 is a GFP dropout vector for constructing dsRNA expression plasmids using Golden Gate assembly. *E. coli* transformed with pBTK800 grow into colonies that are visibly fluorescent under blue light (left). Successful cloning replaces the GFP cassette in pBTK800 with the target sequence. Nearly all *E. coli* colonies are non-fluorescent after transformation with a Golden Gate assembly reaction combining pBTK800 and the *def1* dsRNA target sequence (right). One of these would be picked to verify that it contained the properly assembled dsRNA expression plasmid.

##### Design dsRNA expression plasmid

2. Design primers that amplify a portion of the gene of interest’s coding sequence (the “knockdown target”) (Fig. 2B). Common practice is for this target to be at least 400 bp long, if possible. This length is not a hard requirement, and shorter targets may be successful in silencing shorter genes. Increasing the target length, up to the length of the full gene, may improve target knockdown. Targets can be designed so that they either (A) bridge an exon-intron junction or (B) are entirely contained within an exon. This choice determines how the targets are PCR amplified in step 7. As part of this protocol, we show our design for the knockdown of two bee genes: *insulin receptor 1* (*inR1*, as previously reported^18^) and *defensin-1* (*def1*) (Fig. 2). (1 hour)

3. Design one or two qPCR primer pairs used to assess gene knockdown at the end of the procedure. These primers will not be used until step 53, but designing them beforehand at this early step is important to ensure the amplified products of these primer pairs do not overlap with the targeted region of the gene that will be expressed from the symbiont as dsRNA. Use the IDT qPCR Primer Design Tool (www.idtdna.com/qpcr/design) with default settings. To ensure good efficiency, these primers should amplify a small target (∼100 bp) and have comparable GC contents. We use a target annealing temperature of 60 °C to match the qPCR primers previously designed for the reference gene *rps18*^18^. At this point, one should also order qPCR primers for reference bee genes such as *rps18* and *gapdh*^60,61^. Primer sequences for using qPCR to monitor knockdown of the two example bee genes (*inR1* and *def1*), as well as for reference genes (*rps18* and *gadph*), are shown below. (30 minutes)

Assess *inR1* and *def1* gene expression using the following qPCR primers:

qPCR1_InR1_F: CCCTTCCTCCCTATCTTGTAGTA

qPCR1_InR1_R: GCGAGGAATTGCATGGTTTC

qPCR1_def1_F: TTTATTGTCGGCCTTCTCTTCA

qPCR1_def1_R: TCGGCACGTTCTTCATTCT

qPCR primers for reference genes:

qPCR_GAPDH_F: ACAGACCCGAGTGAATAGATTTG

qPCR_GAPDH_R: CGAACTCAATGGAAGCCCTAA

qPCR_RPS18_F: AGGTGTTGGTCGTCGTTATG qPCR_RPS18_R: CATTCTCCAGCACGCTTATCT

4. Append the following nucleotide sequences to the 5′ end of the target amplification primers designed in step 2. These sequences introduce Golden Gate compatible cut sites (bolded nucleotides below) and overhangs (underlined nucleotides below) for BsaI assembly of the PCR product into the final dsRNA expression vector^16^. (5 minutes)

Forward sequence (5′-3′): GCATCGTCTCATC**GGTCTC**A<u>TATG</u>

Reverse sequence (5′-3′): ATGCCGTCTCA**GGTCTC**A<u>GGAT</u>

5. Order DNA oligonucleotides for assembly and qPCR from IDT or other DNA synthesis service. There is no requirement for special purification. (2 days)

For example, the final primers ordered to build the *inR1* and *def1* knockdown targets are:

InR1_target_F:

GCATCGTCTCATC**GGTCTC**A<u>TATG</u>CCAGATTCCTCACCGTTATGTTTATG

InR1_target_R:

ATGCCGTCTCA**GGTCTC**A<u>GGAT</u>AATCCGAACATAATGAACGAGTTGAG

Def1_target_F:

GCATCGTCTCATC**GGTCTC**A<u>TATG</u>AGAAGAGTAACTTGTGACCTTCTCTC

Def1_target_R:

ATGCCGTCTCA**GGTCTC**A<u>GGAT</u>TGAATCTTCATAATGGCACTTAACCG

##### Amplify dsRNA knockdown sequence

6. Obtain *A. mellifera* DNA. Depending on the target design, the PCR template will be either *A. mellifera* genomic DNA or cDNA. If the desired cDNA target sequence within a gene does not bridge any introns, the researcher may use genomic DNA to amplify the target (A). If the target design bridges an intron sequence, however, the researcher should use cDNA prepared from *A. mellifera* RNA as the template to exclude the intron (B). Alternately, the researcher may order the entire knockdown region as a gBlock (IDT) or similar dsDNA gene fragment. (2 hours)

A(i). Isolate genomic DNA from honey bees using the QIAGEN DNeasy Blood & Tissue Kit according to manufacturer’s instructions. DNA can be isolated from one or more whole bees. Eluted DNA will be used as template for the PCR reaction in step 7.
B(i). Isolate RNA from adult *A. mellifera* using the Zymo Quick-RNA Tissue/Insect Kit according to manufacturer’s instructions. RNA can be isolated from one or more bees, and can be prepared from pulled guts, whole abdomens, bee heads, or whole bees. Use an appropriate body region that expresses your gene transcript at a high level. Eluted RNA will be used as template for a cDNA reaction.
B(ii). Prepare cDNA using Takara 1^st^ strand cDNA Synthesis Kit according to manufacturer’s instructions. Use at least 100ng RNA as template, or more for poorly expressed genes. Either random hexamer or polyT primers can be used as primers for the cDNA reaction. Eluted cDNA will be used as template for the PCR reaction in step 7.

7. Amplify the knockdown target sequence using the constructed primers and appropriate template DNA. Use NEB Q5 Hot Start High-fidelity 2X Master Mix and follow manufacturer’s instructions on reaction setup and thermocycling steps. (5 hours)

8. Verify amplification of a DNA product of the expected size for the knockdown target using agarose gel electrophoresis (Fig. 2C). (1 hour)

**<TROUBLESHOOTING>**

9. Purify the desired DNA product using the QIAGEN QIAquick PCR purification kit. (30 minutes)

10. Quantify the DNA concentration of purified PCR product using a Nanodrop. (10 minutes)

##### Assemble dsRNA expression plasmid via Golden Gate Assembly

11. Prepare for dsRNA expression vector assembly. This reaction can be performed in one of two ways: a “two-part” assembly reaction with plasmid pBTK800 and your PCR product (recommended, procedure A), or a “seven-part” assembly reaction using plasmids pBTK403, pBTK301, pYTK002, pYTK072, pBTK150, pBTK151, and your PCR product (procedure B). pBTK800 is a fluorescent dropout vector described for the first time in this protocol for the simpler and more efficient two-part assembly reaction. This procedure is recommended for a typical user. The seven-part assembly reaction was used previously^18^. It allows components of the plasmid, such as the forward and reverse promoters or the antibiotic resistance marker, to be switched out with other BTK-compatible parts^16^. It may be especially useful for researchers who want more flexibility for designing customized dsRNA expression vectors. (2 days)

A(i). Prepare purified pBTK800 plasmid. Streak from frozen stock onto an LB plate supplemented with spectinomycin. Incubate at 37 °C overnight. All colonies should be visibly fluorescent from expression of GFP. Pick a single fluorescent colony into liquid LB with spectinomycin and grow overnight with shaking in a 37 °C aerobic incubator.
A(ii). Isolate plasmid from stationary phase culture using the PureLink Quick Plasmid Miniprep Kit. pBTK800 has an RSF1010 origin of replication and is a medium copy number plasmid in *E. coli* (∼10-20 copies per cell). Follow kit protocol recommendations for low copy plasmids and use ∼5 mL of overnight culture for the minipreps to purify a sufficiently concentrated sample of this plasmid (≥ 20 ng/µL).
A(iii). Quantify plasmid concentration. Use a Nanodrop to quantify the DNA concentration of purified plasmid. DNA quantification methods based on binding of fluorescent dyes to dsDNA, such as those used by a Qubit Fluorometer, will give incorrect values for the concentration of purified RSF1010 plasmid samples because they contain significant amounts of single-stranded plasmid DNA.
B(i). Prepare purified pBTK403, pBTK301, pYTK002, pYTK072, pBTK150, pBTK151. For each plasmid, streak from frozen stocks onto LB plates supplemented with either chloramphenicol (pBTK301, pYTK002, pYTK072, pBTK150, pBTK151) or spectinomycin (pBTK403). Grow cultures of each strain as described in procedure A, except substituting the appropriate antibiotic. Cells with pBTK403 express RFP and all others are non-fluorescent.
B(ii). Isolate plasmids from stationary phase cultures as described in procedure A. pBTK403 has an RSF1010 origin of replication and should be purified in the same way as pBTK800. All other plasmids are high copy number. For these, sufficient yields can be purified from 1.5 mL of culture using normal procedures.
B(iii). Quantify plasmid concentration. Use a Nanodrop to quantify the DNA concentration of each purified plasmid.

12. Assemble the dsRNA expression vector. Use the NEB Golden Gate Assembly Kit (BsaI-HFv2) mix according to the manufacturer’s instructions to set up a BsaI Golden Gate Assembly reaction. Thermocycle the reaction as recommended by NEB. (1 hour)

##### Transform dsRNA expression plasmid into E. coli cloning strain

13. Transform 1-2 µL of the assembly reaction mixture into NEB 5-alpha Competent *E. coli* (High Efficiency) cells according to manufacturer’s instructions. Incubate cells and recover them for at least one hour before plating dilutions onto LB plates supplemented with spectinomycin. Incubate plates overnight at 37 °C. Colonies containing properly assembled plasmid will be non-fluorescent, while colonies that are fluorescent contain the original, unmodified plasmid (Fig. 2D). Agar plates can be placed over a blue light box to visualize fluorescent colonies. If using approach (A) from step 11, colonies with incorrectly assembled plasmid will exhibit green fluorescence. If using approach (B) from step 11, colonies with incorrectly assembled plasmid will exhibit red fluorescence. (1 day)

**<TROUBLESHOOTING>**

14. Pick at least three non-fluorescent colonies into 5 mL LB with spectinomycin. Incubate at 37 °C in a shaking incubator. (1 day)

15. Isolate assembled dsRNA expression plasmids from 5 mL culture, as described in step 11A(ii). (1 hour)

16. Verify that each assembled dsRNA expression plasmid contains the desired insert using whole plasmid sequencing or targeted Sanger sequencing that covers the cloning junctions between different fragments. Note, Sanger sequencing is not straightforward, given that the insert is flanked on both sides by junctions to identical promoters with sequences that are reverse complements of one another. Use of primers that bind to the vector backbone and are oriented to sequence into the insert may yield unreliable reads under standard sequencing conditions. Three approaches have proven more reliable. The first is to perform Sanger sequencing using primers that bind to the vector backbone under “difficult template conditions” (sometimes referred to as “short hairpin [shRNA] conditions”). Another approach is to perform Sanger sequencing using primers that bind within the sequence cloned from *A. mellifera* for expression as dsRNA, making sure to include primers that will result in sequencing across both junctions to verify insertion. The final alternative approach is to perform PCR reactions that span both junctions on a plasmid sample, and then Sanger sequence these PCR products. Save frozen stocks of transformants with verified inserts by adding glycerol to a final 15% (v/v) concentration to *E. coli* cultures. (2 days)

#### Engineer bee symbiont to express dsRNA

##### Transform dsRNA expression vector into E. coli conjugation strain

17. Several steps are needed to transfer the dsRNA expression vector to *S. alvi* (Fig. 3A). First, prepare an electrocompetent stock of an auxotrophic *E. coli* donor strain, such as Mu-free donor (MFD*pir*) or ST18, according to a standard electrocompetent cell preparation protocol^64^. For the remainder of the protocol, this auxotrophic donor strain will be referred to as MFD*pir*, but ST18 could also be used. ST18, which is available from DSMZ (DSM 22074), is a conjugation-competent *E. coli* strain auxotrophic for 5-aminolevulinic acid^59^. If using ST18, substitute 5-aminolevulinic acid (50 μg/ml)^59^ for DAP. (1 day)

**Figure 3.**
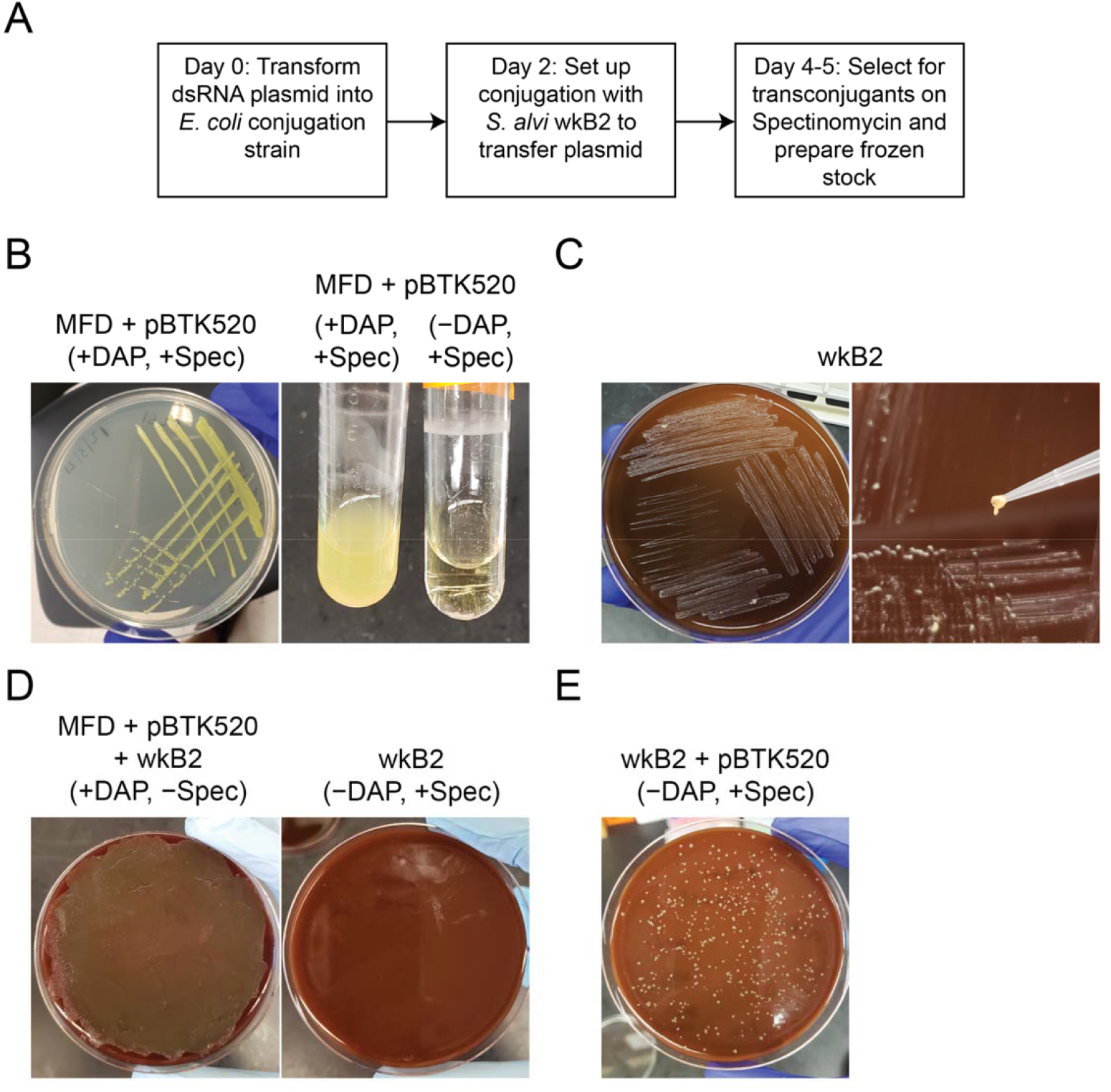
Plasmid conjugation into *S. alvi*. A. Overview of conjugation workflow. B. Conjugation competent *E. coli* donor strain MFD*pir* (MFD) containing a plasmid with spectinomycin resistance, struck out on LB agar +DAP +Spec (left panel). MFD + pBTK520 grown in liquid LB +Spec +/−DAP, showing that MFD is a DAP auxotroph (right panel). The pBTK520 plasmid encodes a constitutively expressed *gfp* that makes *S. alvi* visibly fluorescent^16^. It is used to demonstrate this procedure but is not part of the FUGUES workflow. C. Conjugation recipient *S. alvi* wkB2 strain (wkB2), struck out on Columbia Blood agar (left panel). wkB2 colonies are mucoid in texture and yellowish-white in color when scraped (right panel). D. Conjugation plate containing 9:1 wkB2 recipient:MFD donor; strains grown on +DAP, −Spec plate to facilitate conjugation (left panel). wkB2 control (−DAP, +Spec) does not grow in the presence of Spec (right panel). Both plates are Columbia Blood agar. E. Conjugation selection plate (Columbia Blood agar −DAP, +Spec) showing growth of wkB2 + pBTK520 transconjugants.

**Figure 4.**
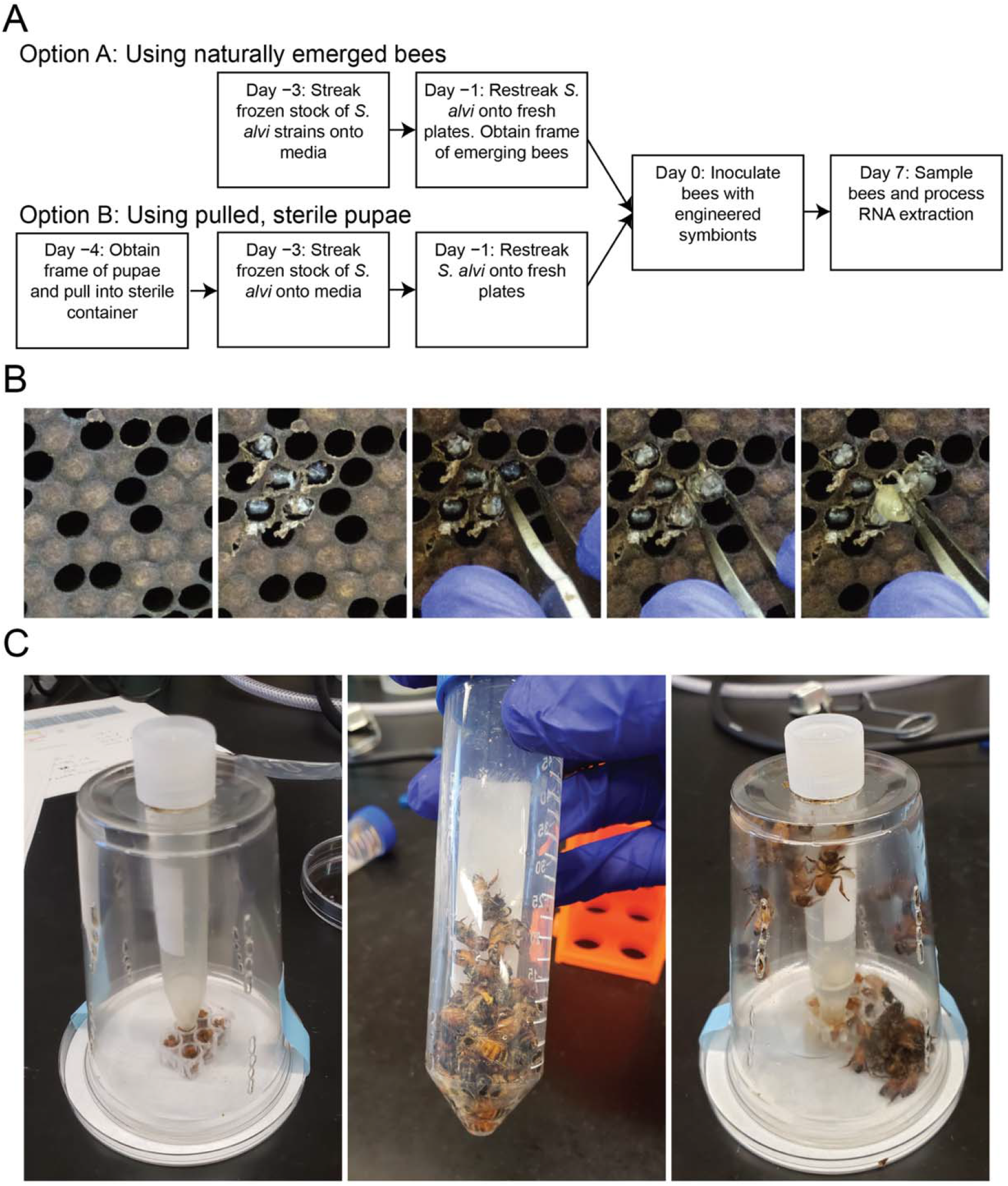
Honey bee inoculation with *S. alvi*. A. Overview of bee inoculation workflow. B. An *A. mellifera* pupa is pulled from a frame (panels move left to right). A frame containing fresh brood is removed from its hive box (left panel) and cells are uncapped (center-left panel) with light scraping. Sterile forceps are carefully inserted into an uncapped cell containing a pupa (center panel), and the pupa is gently pulled (center-right panel) until it is fully removed from its cell (right panel). *C. A. mellifera* adults are inoculated with *S. alvi*. Bee cup cages are set up as described, containing a feeding trough with irradiated pollen and a feeding tube for syrup (left panel). Honey bees are inoculated with *S. alvi* by combining newly emerged adults with a syrup solution containing *S. alvi* in a 50 ml conical tube and shaking to mix (middle panel). Inoculated bees are transferred to a cup cage by removing the feeding tube, inverting the 50 ml tube over the hole in the top of the cup, and replacing the feeding tube (right panel).

Electrocompetent cell preparation protocol^64^:

A. From MFD*pir* frozen stock, streak an LB plate supplemented with DAP and incubate at 37 °C overnight.
B. Pick a single colony into LB media supplemented with DAP. As a control, pick the same colony into LB without DAP. Incubate at 37 °C shaking overnight.
C. Confirm bacterial growth is only in the test tube supplemented with DAP. Dilute the saturated MFD*pir* culture with 10 mL of fresh LB with DAP. Incubate at 37 °C shaking.
D. After 4 hours, begin checking the OD_600_ every 30 minutes until the culture reaches mid-log phase (0.4 – 0.6 OD_600_).
E. Once the culture has reached mid-log phase, transfer the culture to a 15 ml conical tube, centrifuge for 5 minutes at 3824 × g, and remove the supernatant.
F. Add 10 ml cold 10% glycerol to the pellet to wash the cells, keeping cells at 4 °C for the remainder of the protocol (keep cells on ice and use a refrigerated centrifuge cooled to 4 °C). Vortex to mix, centrifuge for 3.5 minutes at 3824 × g, and remove the supernatant.
G. Wash in 10% glycerol, as above, three additional times.
H. Add 100 µl of 10% glycerol to the pellet and pipet to resuspend.
I. Dispense electrocompetent cells into 50 µl aliquots, and freeze at −80 °C if not using immediately for electroporation.

18. Electroporate 1 µL of purified dsRNA knockdown plasmid into electrocompetent MFD*pir* cells. Use the standard *E. coli* setting on the Bio-Rad electroporator compatible with 0.1 cm cuvettes (V = 1.8 kV). (20 minutes)

If this is a researcher’s first time performing this protocol, they must construct strains of *S. alvi* with the negative control plasmids pNR and pDS-GFP. We also recommend that new users follow this protocol with at least one of the plasmids pDS-Def1 and pDS-InR1, which serve as optional positive controls for achieving knockdown of a targeted bee gene. *E. coli* cells containing each of these control plasmids can be obtained from Addgene. Steps 13-31 should be followed to move control plasmids into *S. alvi* in parallel with the target gene plasmids.

19. Recover electroporated MFD*pir* in SOC medium supplemented with DAP for 1 hour, shaking at 37 °C. (1 hour)

20. Plate dilutions of transformed MFD*pir* onto LB plates supplemented with DAP and spectinomycin. Plate a non-transformed control. Incubate overnight and look for colonies the next day (Fig. 3B). (1 day)

**<TROUBLESHOOTING>**

### *Conjugate dsRNA expression plasmid into* S. alvi

21. Pick three colonies into liquid LB supplemented with spectinomycin and DAP. As a control, inoculate 3 test tubes of LB supplemented with spectinomycin (but without DAP) using the same colonies. MFD*pir* donors should only grow in the presence of DAP (Fig. 3B). This is an important control to ensure that the transformed MFD*pir* cells are not contaminated prior to the conjugation. (10 minutes)

22. Make frozen stocks of verified MFD*pir* transformants by adding glycerol to 15% (v/v). It is typically not necessary to sequence the plasmid again at this stage. (10 minutes)

23. Prepare for conjugation into *S. alvi* wkB2. Streak from frozen *S. alvi* stock onto a Columbia Blood agar plate and incubate for 48 hours at 35 °C under 5% CO_2_ (Fig. 3C). Grow the donor strain (MFD*pir* with dsRNA expression plasmid) in LB media supplemented with spectinomycin and DAP. (2 days)

24. Scrape a loopful of wkB2 (Fig. 3C; Video S1) into PBS in a 1.5 mL tube. Transfer 1 mL of MFD*pir* donor culture into a 1.5 mL tube. (5 minutes)

25. Centrifuge both tubes at 3824 × g for 5 minutes. Discard supernatants and resuspend pellet from each tube in 1 mL PBS. Resuspending the *S. alvi* pellet will require repeated pipetting, as *S. alvi* forms a dense and sticky aggregation during growth. (5 minutes)

26. Measure the OD_600_ of each tube using a benchtop spectrophotometer. (2 minutes)

27. Combine wkB2 and MFD*pir* at a ratio of 9:1 by OD_600_ to a total volume of 100 µL and plate on Columbia Blood agar plates supplemented with DAP. Incubate this plate overnight up to 24 hours, allowing both MFD*pir* and wkB2 to replicate (Fig. 3D). As a control, spot wkB2 by itself (no donor) (Fig. 3D) and MFD*pir* by itself (no recipient). (1 day)

28. After overnight growth, use a sterile loop to scrape bacterial growth into 1 mL PBS in a 1.5 mL centrifuge tube. Use a pipette to resuspend the mixture and then spin at 3824 × g for 5 minutes. Discard supernatant and add 1 mL PBS. Pipette to resuspend the cell pellet. Repeat spin at 3824 × g for 5 minutes, discard supernatant, and resuspend in 1 mL PBS as before. This wash is critical for removing residual DAP that could lead to continued MFDpir growth. (10 minutes)

29. Plate dilutions (50 µL, 10 µL, 1 µL) onto Columbia Blood agar plates supplemented with spectinomycin, but without DAP. Incubate plates at 35 °C under 5% CO_2_ for 48 hours. (2 days)

30. Successful transconjugant colonies should be visible after 48-72 hours (Fig. 3E). Streak 1-3 colonies onto fresh Columbia Blood agar plates supplemented with spectinomycin. Incubate for 48 hours at 35 °C under 5% CO_2_. (2 days) **<TROUBLESHOOTING>**

31. Scrape transconjugant wkB2 growth directly into a cryovial with 15% (v/v) glycerol and store at −80 °C. Mix well by pipetting to break up adhesive clumps of *S. alvi*. These stocks will be used for inoculating honey bees. These transconjugants can be confirmed to be pure *S. alvi* cultures by performing 16S rRNA sequencing to ensure no unexpected contaminants have been introduced during the conjugation process. (10 minutes)

#### Colonize bees with engineered symbiont

##### Revive S. alvi to prepare for inoculating bees

32. Several coordinated steps must be carried out to colonize bees with the engineered symbiont (Fig. 4A). Three days prior to the planned inoculation day, streak out colonies from frozen stocks of *S. alvi* wkB2 + dsRNA expression vector (constructed in Step 31), as well as control strains wkB2 + pNR (empty plasmid) and wkB2 + pDS-GFP (off target control). All strains should be struck on Columbia Blood agar plates supplemented with spectinomycin. Incubate strains for 48 hours at 35 °C under 5% CO_2_. Inspect for normal growth or contaminants. (2 days)

33. After 48 hours (one day before planned inoculations), restreak a loopful of bacterial of each strain onto a Columbia Blood agar plate supplemented with spectinomycin. This second passage will be used for inoculation after 24 hours of incubation. (1 day)

##### ollect bee workers from apiary

34. Collect bees according to one of two methods: (1) Allow adult bees to emerge from the frame (1 day) or (2) Pull pupae and allow bees to develop in plastic brood chambers (3-4 days). Method 1 can be performed on the same day as step 33, whereas method 2 should be initiated 3-4 days prior.

###### Method 1

A. Inspect the beehives for a suitable brood frame by gently removing the caps of brood cells with the hive tool and noting the age of pupae. The frame should contain a brood patch with pupae that are ready to emerge, and have dark eyes and sclerotized heads. A frame that contains a small number of bees in a brood patch already chewing through their brood caps works well.
B. Move the frame into the laboratory using an enclosed frame carrier.
C. Incubate the frame in a growth chamber at 35 °C and 20% relative humidity overnight to allow for adult bees to emerge.

###### Method 2

A. Inspect the beehives for pupae 3-4 days earlier than for emerged bees. Look for a brood frame containing brood with dark eyes and partially sclerotized heads, approximately 20-21 days old.
B. Move the frame into the laboratory using an enclosed frame carrier.
C. Once in the lab, use sterile forceps to carefully remove the cell caps from a patch of brood (Fig. 4B).
D. Use sterile forceps to remove individual pupae (Fig. 4B) and place them into a sterile dish provisioned with sterile pollen and 1:1 sucrose:water solution. Flame sterilize forceps periodically to ensure no cross-contamination.
E. Place the dish with pupae into a growth chamber at 35 °C and 20% relative humidity, and allow pupae to finish development over the next 3-5 days. Once pupae have transitioned into adult emergers and are moving around the dish, they are ready for inoculation.

##### Prepare engineered symbiont inoculant

*35. S. alvi* strains are ready for inoculation into bees after 24 hours of growth on agar plates. On the day of inoculation, scrape several loopfuls of bacteria from each agar plate into 1 mL PBS. Use a 1 mL pipette to resuspend the bacteria (to break down biofilm and facilitate dilutions). Spin in a tabletop centrifuge at 3824 × g for 5 mins. Remove PBS and replace with 1 mL fresh PBS. Pipette up and down to resuspend bacteria. (10 minutes)

36. Prepare a 1:10 dilution of the bacterial suspension in PBS and read OD_600_ using a spectrophotometer. Dilute this mixture to an OD_600_ of 0.1. Add 200 µL diluted bacterial suspension to 800 µL of a filter-sterilized 1:1 sucrose:water sterilized solution and mix well by vortexing. The 1 mL of solution prepared in this way is suitable for inoculating 20-30 bees. The precise OD_600_ used to inoculate bees is flexible, as honey bees inoculated with a wide range of *S. alvi* concentrations reliably colonize to the same carrying capacity^18^. (20 minutes)

##### *Inoculate bees with* S. alvi

37. Using forceps or a gentle finger grip, sort bees into groups of 20-30 and place them into sterile 50 mL conical tubes. Newly emerged adult bees (< 24 hours post emergence) can neither sting nor fly, so this sorting can be done without anesthetizing bees. Microbiota-free bees that are not as precisely age controlled should be chilled briefly at 4 °C before sorting. (1 hour)

38. Add 1 mL of inoculation solution (prepared in step 36 above) to each conical tube with bees, close the tube, and gently shake for 10-15 seconds (Fig. 4C). This will coat the bees in the sugary inoculation solution. Allow 2-3 minutes for the bees to begin cleaning each other off. Repeat steps 37-38 until all experimental groups have been colonized. (1 hour)

39. Transfer inoculated bees into cup cages prepared with sterile pollen (Fig. 4C) and place into a humidity-controlled incubator. Allow 30 minutes for bees to entirely clean each other off. The top of the cup cage can be covered with a small petri dish or tape. (30 minutes)

40. After 30 minutes, introduce a feeder containing sucrose solution supplemented with 60 µg/mL spectinomycin. Place the feeder on the pollen trough to catch excess liquid. This feeder should be replaced every 72 hours with fresh sucrose solution and antibiotic. Ensure the feeder tube is neither clogged nor draining too quickly. The feeding solution should be pre-warmed in the incubator to prevent it from excessively leaking when added to cup cages. (30 minutes)

41. Maintain bees in an incubator at 35 °C and 20% relative humidity for 5-21 days. The timing will depend on when knockdown of the gene is expected to have an effect on a bee phenotype of interest, which may need to be empirically determined by a researcher. Check daily for any deceased bees and remove them from cup cages. *S. alvi* should fully colonize the bee gut 3-5 days after inoculation. Bees that have been inoculated with the engineered symbionts will generally survive in lab for at least 2-3 weeks. (5-21 days)

42. (Optional) After incubating bees with engineered symbiont for 5 days, validate colonization. Dissect guts from inoculated bees, homogenize the tissue in a 1.5 ml tube (resuspend in 500 ul PBS and grind with a pestle), plate dilutions onto Columbia Blood agar plates containing spectinomycin, incubate at 35 °C, 5% CO_2_ for 2-3 days^18^, and record CFUs. We typically observe that engineered *S. alvi* numbers climb to a stable level of ∼5 × 10^7^ CFU per bee by 5 days after colonization^18^. (2-3 days)

**<TROUBLESHOOTING>**

#### Validate gene knockdown

##### Sample bees

43. The final steps in this protocol are used to test for knockdown of the target gene (Fig. 5A). After a minimum of 5 days, sample bees from the experimental and control groups (positive control: *inR1* knockdown or *def1* knockdown; negative control: *gfp* mock knockdown). Sample bees by immobilizing them using (A) CO_2_ or (B) chilling to 4 °C.

**Figure 5.**
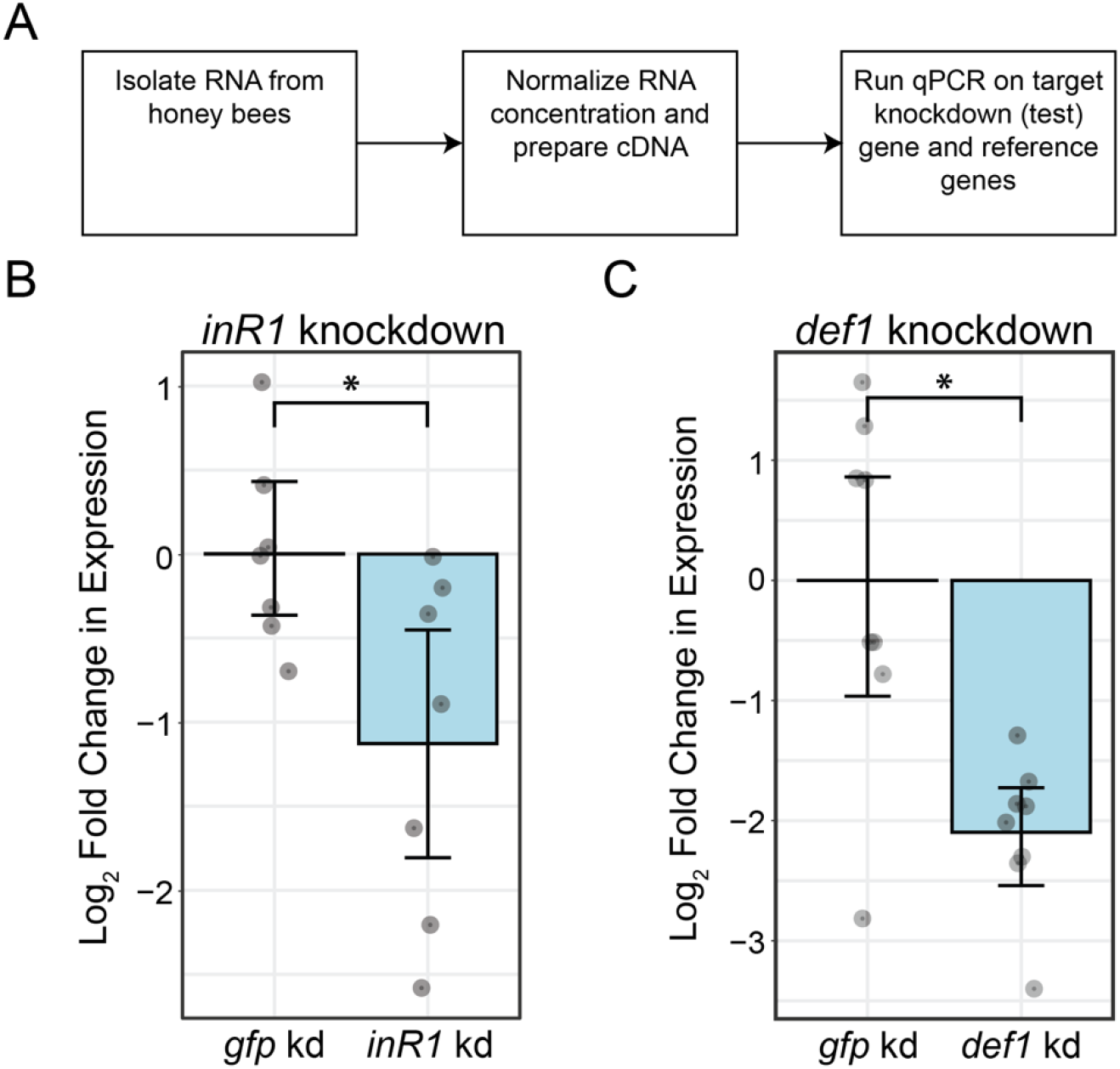
Validation of gene knockdown. A. Overview of gene knockdown validation workflow. B. Fold expression plot showing decreased *inR1* gene expression in bees treated with the *inR1* knockdown symbiont strain. qPCR was performed on cDNA derived from bee guts that were dissected from bees colonized with *S. alvi* expressing dsRNA sequences matching either *inR1* (*inR1* kd) or *gfp* (*gfp* kd; off-target control) for 5 days. Expression levels of *inR1* (the test gene) were normalized to *rps18* and *gapdh* (reference genes). Expression of *inR1* was significantly lower in bees treated with the on-target dsRNA versus the off-target control (Mann-Whitney *U* test, *p* = 0.038). N = 7 for both *inR1* and *gfp*. Data for *inR1* is from a published study^18^. C. Fold expression plot showing decreased *def1* gene expression in bees treated with the *def1* knockdown symbiont strain. qPCR was performed on cDNA derived from bee abdomens that were dissected from bees colonized with *S. alvi* expressing dsRNA sequences matching either *def1* (*def1* kd) or *gfp* (*gfp* kd; off-target control) for 7 days. Expression levels of *def1* (the test gene) were normalized to *rps18* and *gapdh* (reference genes). Expression of *def1* was significantly lower in bees treated with the on-target dsRNA versus the off-target control (Mann-Whitney *U* test, *p* = 0.0070). N = 8 for both *def1* and *gfp*.

A. Place entire cup cage in a plastic bag and introduce CO_2_ until bees stop moving. Bees will be paralyzed within 30 seconds of CO_2_ exposure, and remain paralyzed for 1-2 minutes.
B. Place entire cup cage in a refrigerated environment and wait until bees stop moving. This may take 30 minutes to 1 hour depending on the number of bees in the cup cage.

Working quickly, remove 8-12 immobilized bees from the cup cage and place them in either a 5 mL conical tube for long term storage at −80 °C or directly on a petri dish over ice for immediate processing. Bees can be stored at −80 °C for at least 6 months and then processed for RNA isolation or can be processed immediately. (1-2 hours)

44. (Optional) Dissect the appropriate bee tissue that expresses the gene of interest prior to RNA extraction. Sever the head or abdomen using a clean razor blade for genes expressed in brain tissue or the fat body, respectively. Whole guts are recommended for initial tests using the positive and negative controls, as robust gene knockdown is observed in the gut^18^. Tissue used for experimental groups and related controls will depend on the researcher’s experiment. If not dissecting guts, proceed to the next step. To dissect whole guts, ensure an individual bee is immobile on a petri dish placed above a bucket of ice. If using samples previously frozen at −80 °C, allow bees to thaw on ice before dissection. (1 hour)

A. Grasp the bee thorax using a pair of flat tipped forceps in one hand and grasp the last tergite before the bee stinger with a pair of sterile point-tipped forceps.
B. While holding the thorax in place, smoothly pull the point-tipped forceps away from the thorax. As the tip of the abdomen separates from the rest of the abdomen, the entire unpunctured gut should remain attached to the stinger.
C. Place the whole gut into the prepared tubes.
D. Flame or alcohol sterilize forceps and repeat this procedure for all bee samples.

##### Isolate total RNA from bees

45. Process all samples for RNA extraction by first placing the bee tissue in a 1.5 ml centrifuge tube and manually homogenizing with a pestle. Then, follow the manufacturer’s instructions for RNA extraction using the Zymo Quick-RNA Tissue/Insect Kit. For the initial bead beating step, processing two times for 30 seconds each on a BioSpec Mini-Beadbeater-96 cell disruptor is sufficient. Perform DNAse treatment clean up on each sample of isolated RNA according to manufacturer’s instructions. (3 hours)

46. Elute total RNA in 20 µL molecular grade water into RNase-free microcentrifuge tubes. (5 minutes)

47. Quantify concentration of RNA using Nanodrop. Sample purity can be assessed from the 260/280 ratio, with a value of 2.0 considered to be pure^65^. (5 minutes)

48. Freeze RNA at −80 °C for storage or use immediately for cDNA synthesis. (5 minutes)

##### Generate cDNA from total RNA

49. Generate cDNA using the Takara 1^st^ strand cDNA Synthesis Kit following the manufacturer’s instructions with a 10 µL reaction volume and random hexamer primers. (30 minutes)

A. Normalize total input RNA for all samples. We recommend at least 100 ng RNA per sample. Samples with very low concentrations of RNA due to improper extraction or RNase contamination should be excluded at this step.
B. All samples from a given experiment that will be compared to each other during qRT-PCR should be processed together to minimize batch effects.

50. Dilute 10 µL of cDNA with 90 µL molecular grade water so that there is a sufficient volume of this template to perform several qRT-PCR reactions on each sample that amplify the relevant reference and test genes. (5 minutes)

##### Perform qRT-PCR for reference and target genes

51. Test qPCR primer efficiency by generating a standard curve. Before using qPCR primers on test samples, ensure the designed qPCR primers have an amplification efficiency close to 100% and that it is approximately equal to the reference genes. Measure efficiency by performing qPCR on a serial 10-fold dilution of a PCR amplified gene product. A single 96-well qPCR plate can run a standard curve for one primer set (Dilutions 10^0^-10^−7^) and a total of 24 samples in triplicate. For large experiments, use a 384-well qPCR plate if possible. (2 hours)

A. Add bee cDNA and qPCR primer sets (for reference and test genes) to a 50 µL PCR reaction using OneTaq 2X Master Mix with Standard Buffer according to the manufacturer’s instructions. 1-2 µL of diluted cDNA should be a sufficient amount of template.
B. Thermocycle the PCR reaction according to the manufacturer’s instructions. Choose an annealing temperature that matches that of your designed primers.
C. Confirm amplification of the gene product occurs at the intended size (∼100 bp) by performing gel electrophoresis on the PCR product.
D. Clean up the PCR amplicon using the QIAGEN QIAquick PCR purification kit according to the manufacturer’s instructions.
E. Elute the purified PCR product in 50 µL molecular grade water.
F. Use the purified PCR product as template for a qPCR reaction with eight 1:10 serial dilutions to cover a large dynamic range of amplification. Each dilution should be run in triplicate. Run qPCR using Bio-Rad 2× SYBR Green Master Mix according to the manufacturer’s instructions.
G. After completion of the qPCR reaction, generate a standard curve using the Ct values for each dilution and the log of the starting DNA concentration.
H. Calculate efficiency using the formula *E* = 10^−(1/slope)^ – 1, where “slope” is the slope of the standard curve generated above^66,67^. Multiply this value by 100 to get % efficiency. For the ΔΔCt comparison method (described below) to be valid, efficiencies for the reference and test genes should be roughly equivalent^68^. If these values differ substantially, use the Pfaffl method for calculating relative changes in gene expression^68^.

52. Perform qPCR on at least 7-8 samples each from the experimental and control groups. Each sample will be used for at least two qPCR reactions, one with primers for a reference gene and one with primers for a test gene. If desired, multiple reference genes can be used^60,61^. We describe using qPCR primers that target reference genes *rps18* and *gapdh* that are provided in step 3. If possible, set up all samples to be compared on a single qPCR plate, to minimize run-to-run variability. If this is not possible, ensure each qPCR plate has samples from every experimental group to minimize batch effects. (2 hours)

A. Prepare dilutions of PCR standards for each primer set. Serially dilute 1 µL of purified PCR product across seven 10-fold serial dilutions, and use pure water for the eighth sample as a negative control. These dilutions and negative control will be on every qPCR plate set up using these primer pairs as a control.
B. Prepare qPCR master mix on ice. For each sample prepare triplicate 10 µL reactions that include:
  - 5 µL Bio-Rad 2× SYBR Green Master Mix
  - 0.5 µL each qPCR primer (10 µM concentration)
  - 3 µL Molecular grade water
  - 1 µL template (diluted cDNA reaction from step 50)
  For consistency, we recommended preparing a master mix that contains enough of all of these components (×1.1 for pipetting error) for every sample, except for the unique template. Gently pipette to mix.
C. Dispense 9 µL of master mix into every well of a qPCR plate. The plate should be set up on ice. For consistency and easy setup, we recommend using multichannel pipettes.
D. Dispense 1 µL of each standard prepared in step 51A in triplicate.
E. Dispense 1 µL of each experimental sample into a well in triplicate.
F. Use a centrifuge to collect liquid at the bottom of each well.
G. Apply an optically clear plastic cover to the qPCR plate.
H. Load the qPCR plate into a pre-heated qPCR machine (Mastercycler ep realplex Real-time PCR System). Perform the thermocycling reaction according to default parameters, according to manufacturer’s instructions.
I. At the end of the run, export the raw Ct values (which will be used to determine changes in gene expression, with the ΔΔCt method).

##### Calculate fold expression and run statistical analyses

53. Calculate ΔΔCt using previously described methods^62,63^. This will require solving for two Ct values, one for the reference gene and one for the test gene. (10 minutes) Briefly:

A. Inspect the standard curves to ensure efficiency for the primer pair is as expected.
B. Calculate the mean Ct value for the reference gene and the mean Ct value for the test gene for the control samples.
C. Calculate the mean Ct value for the reference gene and the mean Ct value for the test gene for the experimental samples.
D. Calculate ΔΔCt for the experimental and control samples by subtracting mean control sample Ct from calculated Ct values.

Alternatively, R packages such as `pcr` or `qpcR` can be used to calculate fold changes in expression^62,69^.

54. Convert ΔΔCt values to relative expression using the following equation^62^: Relative expression = 2^−^ΔΔ^CT^. Optionally, further log_2_ transform the relative expression data. (5 minutes)

55. Plot relative expression values (untransformed or log_2_-transformed) using graphing software to visualize changes in gene expression (compared to the reference gene) in experimental versus control samples (Fig. 5B-C). (20 minutes)

56. Analyze the change in gene expression in the experimental group compared to the control group to determine if gene expression knockdown is statistically significant. First, check if data points within each group are normally distributed using D’Agostino’s *K*^2^ test. If they are, then an unpaired *t*-test can be used to test for a significant difference. If the data are not normally distributed, then the nonparametric Mann-Whitney *U* test can be used. (5 minutes)

**<TROUBLESHOOTING>**

### Timing

Clone dsRNA expression vector (Steps 1-16, 7 days)

Engineer bee symbiont to express dsRNA (Steps 17-31, 9 days)

Colonize bees with engineered symbiont (Steps 32-42, 10-27 days)

Validate gene knockdown (Steps 43-56, 1-2 days)

### Troubleshooting

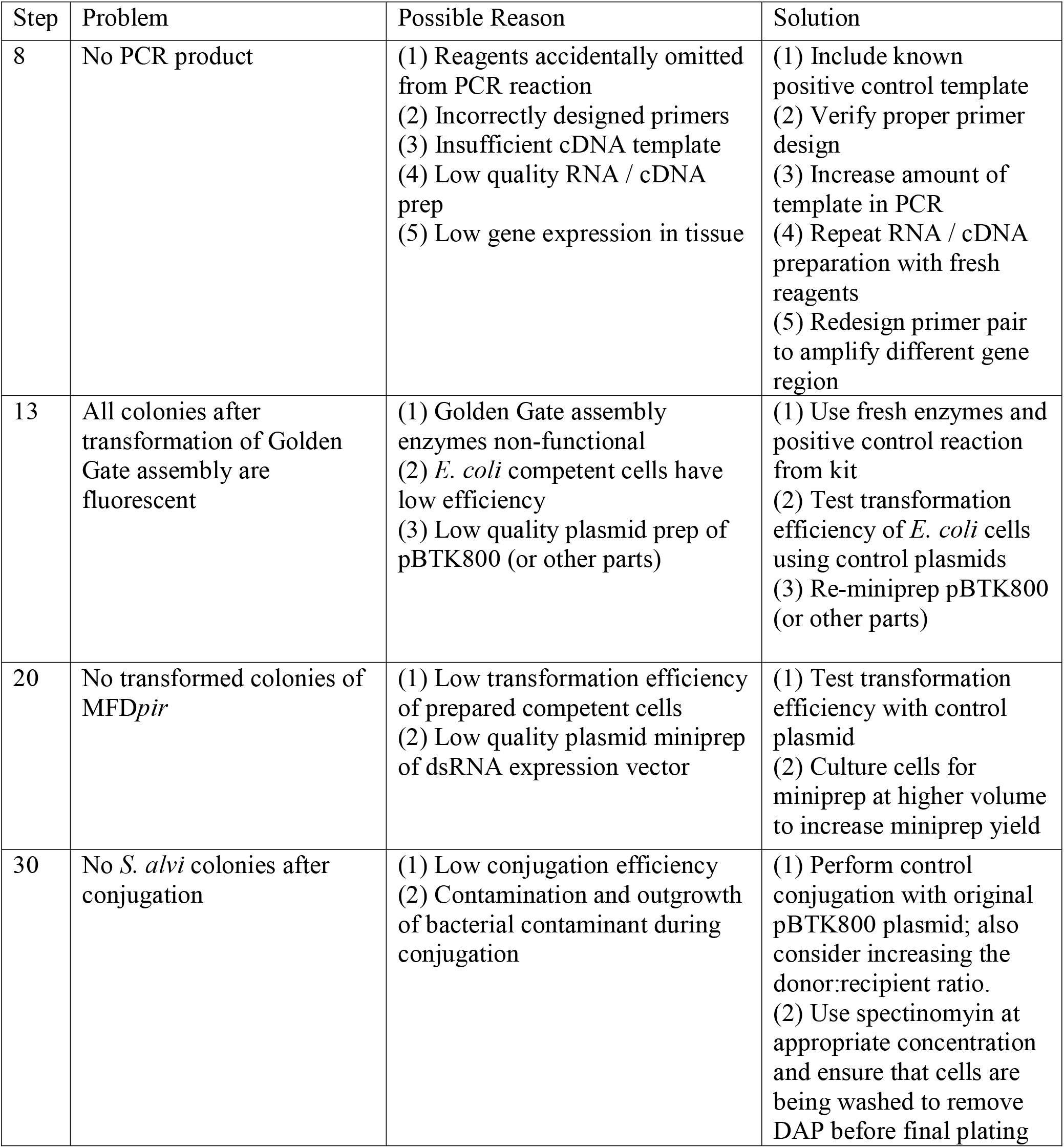
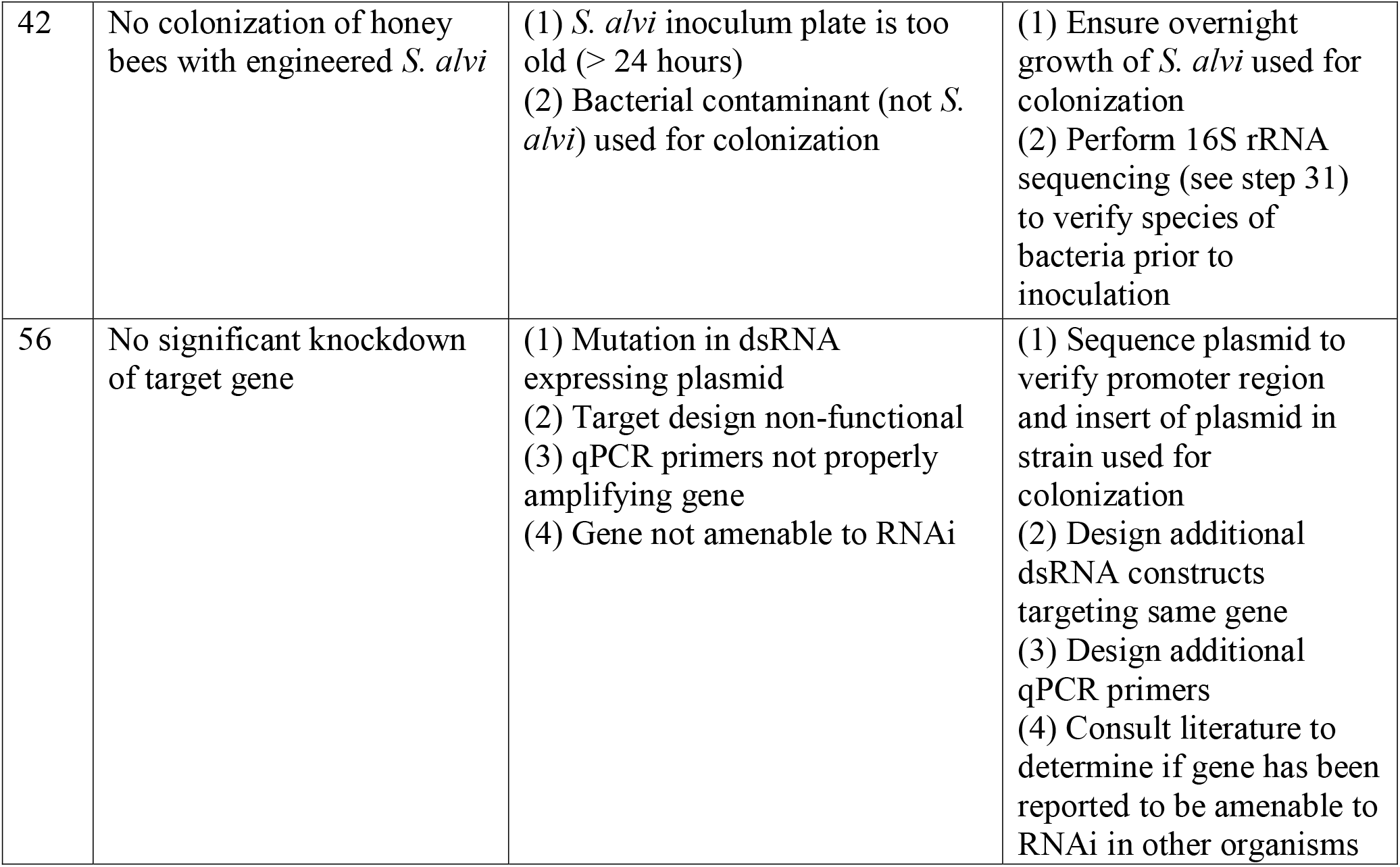

| Step | Problem | Possible Reason | Solution |
| --- | --- | --- | --- |
| 8 | No PCR product | (1) Reagents accidentally omitted from PCR reaction<br>(2) Incorrectly designed primers<br>(3) Insufficient cDNA template<br>(4) Low quality RNA / cDNA prep<br>(5) Low gene expression in tissue | (1) Include known positive control template<br>(2) Verify proper primer design<br>(3) Increase amount of template in PCR<br>(4) Repeat RNA / cDNA preparation with fresh reagents<br>(5) Redesign primer pair to amplify different gene region |
| 13 | All colonies after transformation of Golden Gate assembly are fluorescent | (1) Golden Gate assembly enzymes non-functional<br>(2) *E. coli* competent cells have low efficiency<br>(3) Low quality plasmid prep of pBTK800 (or other parts) | (1) Use fresh enzymes and positive control reaction from kit<br>(2) Test transformation efficiency of *E. coli* cells using control plasmids<br>(3) Re-miniprep pBTK800 (or other parts) |
| 20 | No transformed colonies of MFD<sub>pir</sub> | (1) Low transformation efficiency of prepared competent cells<br>(2) Low quality plasmid miniprep of dsRNA expression vector | (1) Test transformation efficiency with control plasmid<br>(2) Culture cells for miniprep at higher volume to increase miniprep yield |
| 30 | No *S. alvi* colonies after conjugation | (1) Low conjugation efficiency<br>(2) Contamination and outgrowth of bacterial contaminant during conjugation | (1) Perform control conjugation with original pBTK800 plasmid; also consider increasing the donor:recipient ratio.<br>(2) Use spectinomycin at appropriate concentration and ensure that cells are being washed to remove DAP before final plating |
| 42 | No colonization of honey bees with engineered *S. alvi* | (1) *S. alvi* inoculum plate is too old (> 24 hours)<br>(2) Bacterial contaminant (not *S. alvi*) used for colonization | (1) Ensure overnight growth of *S. alvi* used for colonization<br>(2) Perform 16S rRNA sequencing (see step 31) to verify species of bacteria prior to inoculation |
| 56 | No significant knockdown of target gene | (1) Mutation in dsRNA expressing plasmid<br>(2) Target design non-functional<br>(3) qPCR primers not properly amplifying gene<br>(4) Gene not amenable to RNAi | (1) Sequence plasmid to verify promoter region and insert of plasmid in strain used for colonization<br>(2) Design additional dsRNA constructs targeting same gene<br>(3) Design additional qPCR primers<br>(4) Consult literature to determine if gene has been reported to be amenable to RNAi in other organisms |

### Anticipated results

This honey bee FUGUES protocol yielded 50% to 75% knockdown of gene expression for the positive controls targeting the *inR1* and *def1* genes (Fig. 5B). Knockdown of *inR1* by this amount affected bee phenotypes, including behavior and weight gain^18^. The efficiency of gene knockdown by RNAi can vary substantially depending on target specific effects (such as native gene expression levels and tissue-dependent expression) and design of the dsRNA^29,40,41^. The qPCR procedure described here is the most direct method to confirm knockdown of gene expression when testing the FUGUES approach on a new gene. Once significant knockdown is established, researchers can move on to study whether there are any effects on organismal phenotypes to provide direct evidence that a gene is involved in a biological process of interest.

## Acknowledgements

We thank Nancy Moran for providing invaluable guidance and resources during the development of this protocol. We also thank Benjamin Jack for his help in graphic creation (figure 1). Development of this protocol was supported by funding from the National Science Foundation (IOS-2103208) and the Defense Advanced Research Projects Agency (HR0011-15-C0095).

## Author contributions

Conceptualization, S.P.L. and J.E.B.; Methodology, S.P.L., J.E.P., and J.E.B.; Investigation, P.J.L., S.P.L., J.E.P., R.D.H.; Writing, P.J.L., S.P.L., and J.E.B.; Editing, P.J.L., S.P.L., J.E.P., R.D.H., and J.E.B.

## Competing interests

S.P.L. and J.E.B. have a pending patent (US-20190015528-A1) on the use of engineered symbionts to improve bee health. J.E.B. is the owner of Evolvomics LLC.

